# Single cell gene fusion detection by scFusion

**DOI:** 10.1101/2020.12.27.424506

**Authors:** Zijie Jin, Wenjian Huang, Ning Shen, Juan Li, Xiaochen Wang, Peter J. Park, Ruibin Xi

## Abstract

Gene fusions are widespread in tumor cells and can play important roles in tumor initiation and progression. Using full length single cell RNA sequencing (scRNA-seq), gene fusions can now be detected at single cell level. However, scRNA-seq data has a high noise level and contains various technical artefacts that can lead to spurous fusion discoveries. Here we present a computational tool, scFusion, for gene fusion detection based on scRNA-seq. scFusion can efficiently and sensitively detect fusions with a low false discovery rate. In a T cell data, scFusion detected the invariant TCR gene recombinations in Mucosal-associated invariant T cells that many methods developed for bulk-data failed to detect. In a multiple myeloma data, scFusion detected the known recurrent fusion *IgH-WHSC1*, which was associated with overexpression of the *WHSC1* oncogene.

## Introduction

Gene fusions are hybrid genes by joining of two independent genes, often resulting from structural rearrangements such as deletions and translocations^1^. In cancer cells, many gene fusions are driver mutations that play important roles in the carcinogenesis process. Well-known examples include the *BCR–ABL1* fusion in chronic myeloid leukemia^2^, the *TMPRSS2– ERG* fusion in prostate cancer^3^ and the *ALK* fusions in lung cancer^4^. Some gene fusions are strongly correlated with tumor subtypes and are often used as diagnostic markers for malignancy. Gene fusions are also important targets of cancer drugs. A number of such drugs have been approved by the US Food and Drug Administration such as imatinib targeting *BCR-ABL1*^5^, crizotinib targeting *ALK* fusions^6^ and the recently approved TRK inhibitor larotrectinib targeting *NTRK* fusions^7^.

RNA sequencing (RNA-seq) provides an accurate and unbiased platform for gene fusion detection. Gene fusions generate chimeric reads that cover junctions of independent partner genes. After mapping, these chimeric reads will be split-aligned to the partner genes and gene fusions can be detected. The development of full length single cell RNA sequencing (scRNA-seq) ^8-10^ unprecedentedly enables the detection and investigation of gene fusions at single cell level. Fusion detection at single cell level could allow identification of cell subtypes or subclones that play unique roles in the complex biological systems and/or have unique phenotypic, genotypic and transcriptomic features. However, although many computational fusion detection tools have been developed for bulk RNA-seq data^11-19^, fusion detection based on scRNA-seq is still challenging: (1) The heavy amplification step of scRNA-seq may generate artifacial chimeric reads and lead to false positive gene fusion discoveries; (2) Many single cells can have similar transcriptomic features and the power for detecting fusions shared among multiple single cells can be improved if all single cells are integratively analyzed. (3) Current single cell data often contain thousands or even more single cells. Although one cell’s data is generally smaller than a bulk RNA-seq data, collectively, scRNA-seq data is large. Fusion detection methods should be optimized for analyzing large single cell data.

In this paper, we developed a gene fusion detection algorithm named scFusion based on the full length scRNA-seq data. scFusion employs a statistical model and a deep learning model to control for the false positives.To assess the performance of scFusion, we first simulated single cell data on the basis of real scRNA-seq data and found that scFusion achieved high sensitivity while maintaining a low false discovery rate (FDR). Next, we experimentally added gene fusions to single cells and validated that scFusion could successfully detect these spike-in fusions. Finally, we applied scFusion to two publicly available scRNA-seq data and showed that scFusion identified cell subtypes closely associated with gene fusions.

## Results

### scFusion is a method designed to detect gene fusion from scRNA-seq data

scFusion takes input as reads mapped by STAR^20^ (Fig. 1). Uniquely split-mapped reads and discordant reads mapped to different genes are identified and clustered to obtain candidate gene fusion list. After filtering fusion candidates in pseudogenes, long noncoding RNAs (lncRNAs), genes without an approved symbol ^21^ (such as *RP11-475J5*.*6*), and in the intronic regions, we observed well over 10,000 fusion candidates in all datasets tested

**Fig. 1.**
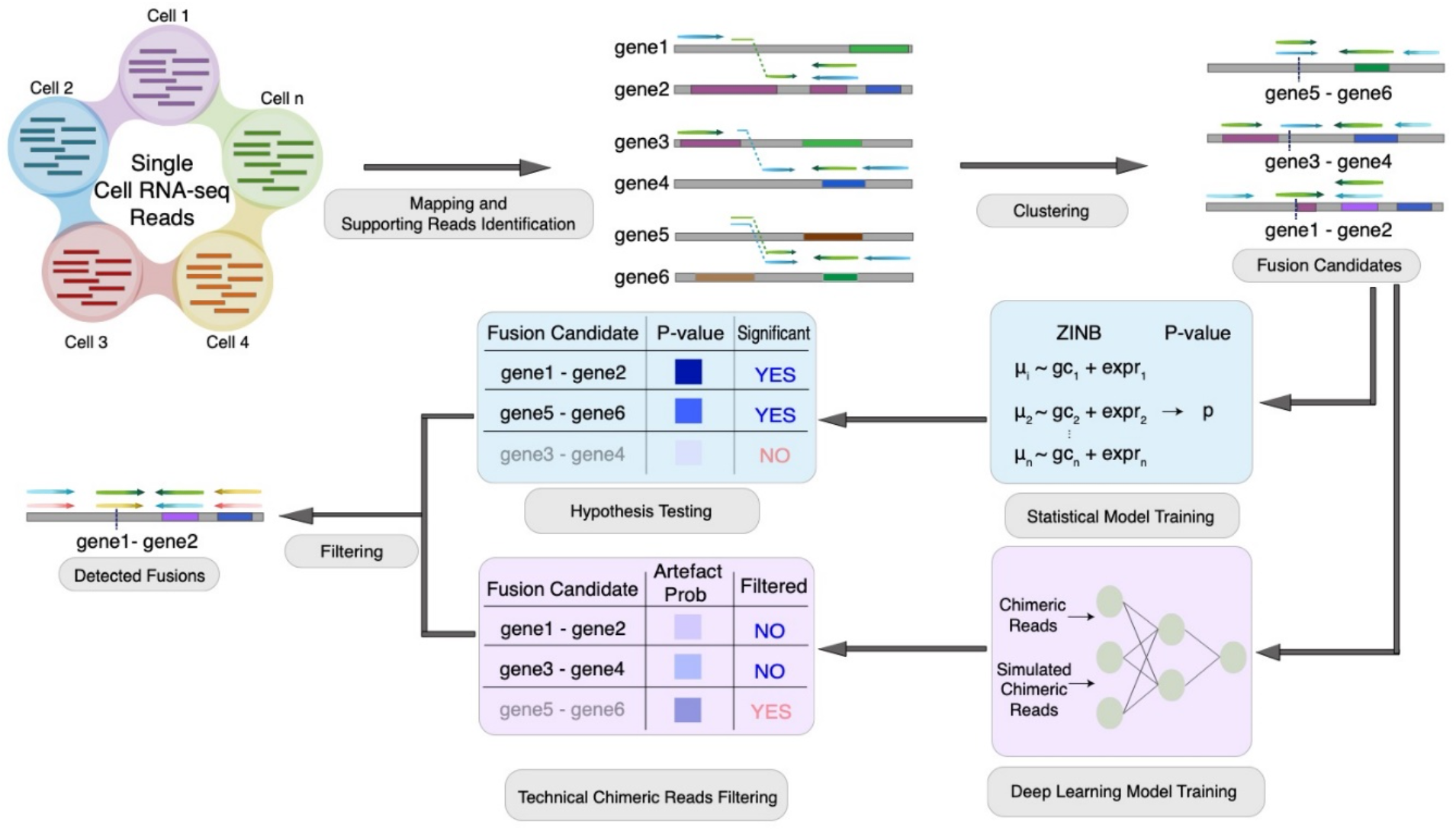
Overview of scFusion for single cell gene fusion detection. The single cell RNA-seq reads are mapped and supporting reads are identified and clustered to obtain the fusion candidates list. Given the candidate information, a ZINB-based statistical model and a deep learning model are trained to filter the potential false positives.

(Extended Data Fig. 1). When considering only candidates found in at least two cells, there were still thousands of fusion candidates, which were mainly false positives. For example, the T cells of a hepatocarcinoma scRNA-seq dataset^22^ had ∼ 3,000 fusion candidates found in at least two cells (excluding candidates involving T cell receptor related genes). Since these T cells are normal cells, true fusions should be very rare in this data set. To control for false discoveries, scFusion independently applies a statistical model and a deep learning model (**Methods**). Finally, scFusion filters out fusion candidates whose number of supporting discordant reads is ten times more than the supporting split-mapped reads and the candidates with a partner gene involving in more than five fusion candidates.

There are two essential assumptions of the statistical model in scFusion: 1) the candidate fusion list only contains a very small fraction of true fusions, and 2) the true fusion transcripts generally should have more supporting cells/reads than background noises with similar characteristics. The distribution of background noises can be estimated using all candidate fusions. True fusions can be identified by comparing with the background distribution. We observed that, for each fusion candidate, the number of supporting chimeric reads in a cell depended on the expression of partner genes and local GC content (Fig. 2a, b). We describe this dependence using a generalized additive model based on the zero inflated negative binomial (ZINB) distribution. After the model parameters are estimated, we perform a statistical test to test fusion candidates not coming from the background noise (**Methods**).

**Fig. 2.**
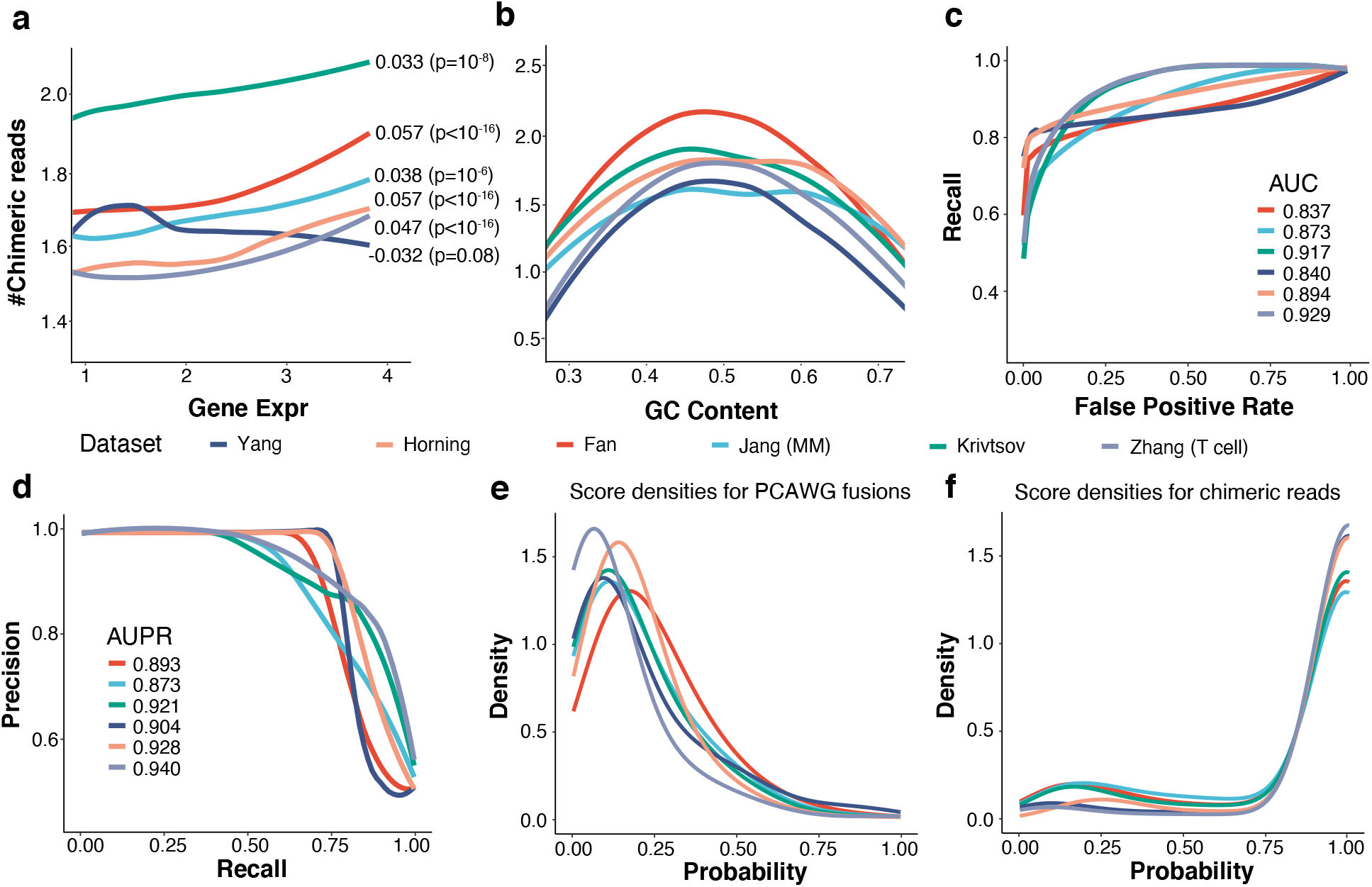
Features of technical chimeric reads. The number of supporting chimeric reads depends on (**a**) the expression of partner genes and (**b**) the local GC content. The Pearson’s Correlations between the number of chimeric reads and the gene expression and the p-values are shown in the figure. The GC content is calculated using sequences near breakpoints (200 bp). (**c**) The ROCs of the bi-LSTM model for different single cell data sets (validation data). The AUCs are also shown. (**d**) The PR curves and their AUPRs. (**e**) The densities of the technical artefact score of gene fusions in the PCAWG study by the bi-LSTM models retrained using six different datasets. (**f**) The densities of the predicted probabilities of chimeric reads. The models are retrained using different datasets.

The statistical model alone is not sufficient to filter out all false fusion candidates since false positive fusion candidates may be induced frequently in single cells during the experimental precedure of cell lysising and sequencing. We observed that the sequences near the junctions of the chimeric reads were enriched with sequences like AAAA and AGGT (Extended Data Fig. 2a), and hypothesized that the technical artefacts might be learned by a machine learning model. We therefore used a bi-directional Long Short Term Memory network (bi-LSTM)^23,24^ to to learn and filter the artefacts (**Methods** and Extended Data Fig. 2b). We set the negative training data as the subsequences from chimeric reads, representing the technical artefacts. Since the number of true fusions are too small for the bi-LSTM training, the positive training data is set as sequences generated by concatenating random pairs of short reads (**Methods**). The bi-LSTM assigns each candidate an artefact score from 0 to 1 and candidates with artefact scores greater than 0.75 are filtered.

We evaluated the performance of the bi-LSTM model using six publicly available scRNA-seq datasets^22,25-29^. The median of the area under curve (AUC) was 0.884 and the median area under the precision-recall (PR) curve (AUPR) was 0.913 (Fig. 2 c,d). Since the bi-LSTM model was not trained using true fusions, true fusions could be filtered by this model. To evaluate this possibility, we applied the bi-LSTM models trained using six public scRNA-seq data to 3500 gene fusions reported in Pan-Cancer Analysis of Whole Genomes (PCAWG) studies^30,31^. The PCAWG fusions can be viewed as gold strandard true fusion candidates. The PCAWG fusions and the chimeric reads had artefact scores centered around 0.1 (Fig. 2e) and 0.95 (Fig. 2f), respectively. More than 90% of the PCAWG fusions had artefact scores smaller than 0.5, and only ∼5% had scores larger than 0.75. These indicated that the bi-LSTM model could effectively filter chimeric artefacts at the expense of filtering a very small portion of true fusions. Further,the bi-LSTM did learn features of chimeric artefacts. Chimeric reads with high artefact scores can be partially attributed to features such as their junction sequences (Extended Data Fig. 2c-g).

### scFusion detects fusions with higher precision in simulation

We first used simulation to evaluate the performance of scFusion and compared with several popular bulk methods including Arriba^18^, STAR-Fusion^19^, FusionCatcher^12^ and EricScript^17^. We used FusionSimulatorToolkit ^19^ to generate simulation data with 150 fusions at various expression levels. Since technical chimeric reads cannot be generated by available simulation tools, we added the chimeric reads from real scRNA-seq data to the simulated data (**Methods**). We set the percentage of chimeric reads to be 1%, similar to those in the real scRNA-seq data (Extended Data Fig. 3a). We varyed single cell numbers (500 and 1,000 cells) and data sizes of each single cell (2, 3 and 4 million reads, similar to those in real data, see Extended Data Fig. 3b). In total, we had six different simulation setups and we generated ten datasets for each setup. Each simulated fusion was randomly added to 20% cells. For bulk methods, we only considered fusions that were reported in at least two cells to reduce their false positives. Compared with bulk methods, scFusion had similar level of recalls but higher levels of precisions and F-scores (Fig. 3 and Extended Data Fig. 4). For example, in the simulation with 1000 single cells and 3 million reads per cell, the precision and F-socre of scFusion are 0.94 and 0.93, respectively, while the precisions of the bulk methods are only 0.13-0.70 and the F-scores are 0.23-0.72. The fusions missed by scFusion are mostly fusions with low expression levels (Extended Data Fig. 5).

**Fig. 3.**
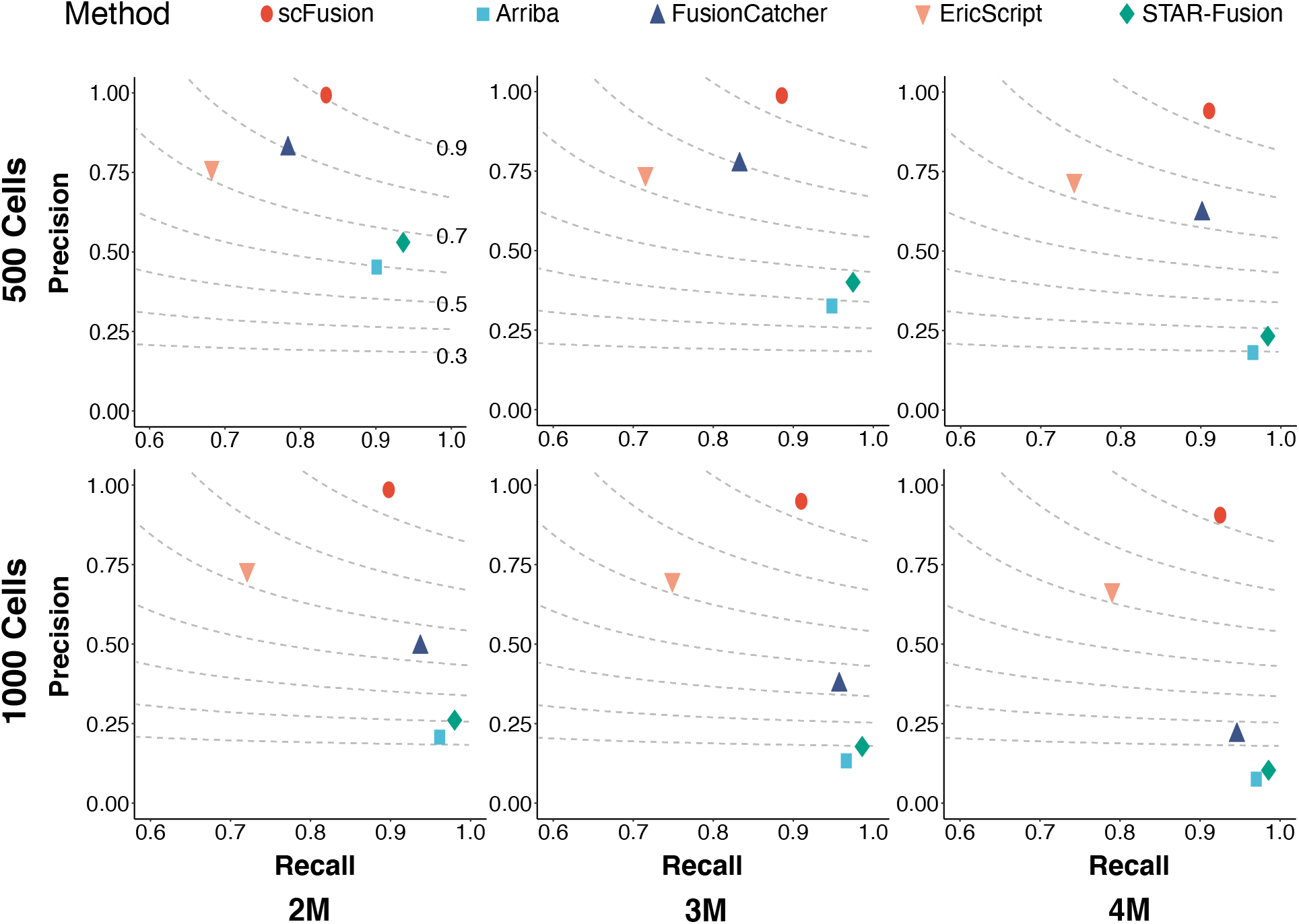
The precisions and recalls of scFusion and four bulk methods in six different simulation setups. The figures in the two rows correspond to simulations with 500 cells and 1000 cells, and the figures in the three columns correspond to simulations with 2 million, 3 million and 4 million reads in each data. The dots in the figures are the means of precisions and recalls of ten simulations in each setup. The dashed lines are the contour lines with constant F-scores (F-scores are marked in the top-left figure).

**Fig. 4.**
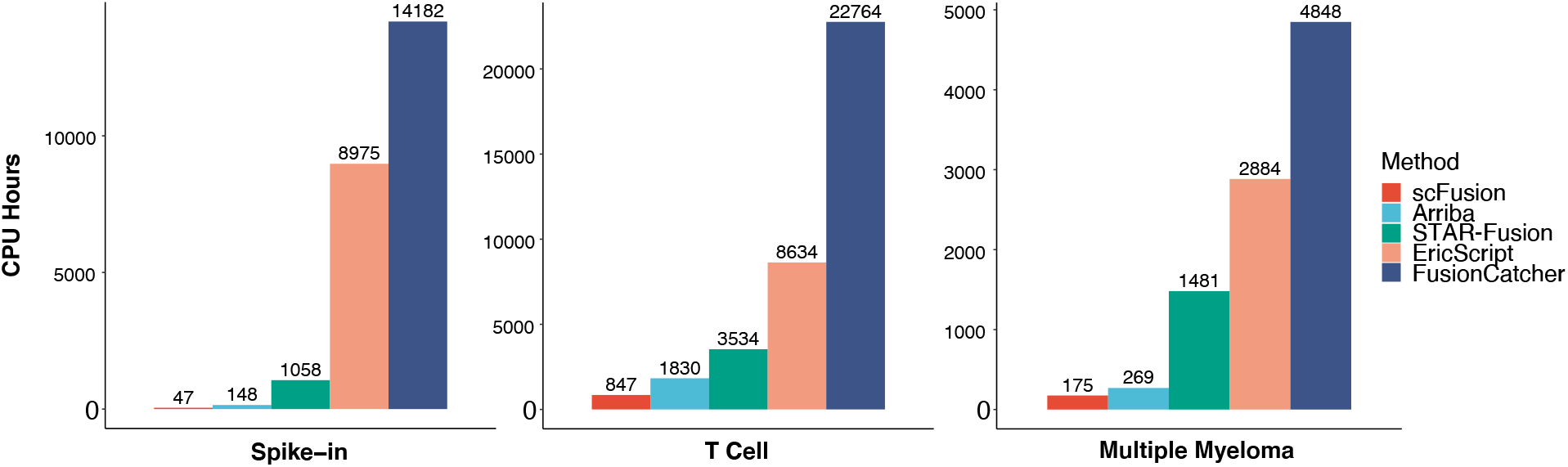
The computational time of the five methods for fusion detection in the three scRNA-seq data. The y-axis is the total CPU hours of each method.

**Fig. 5.**
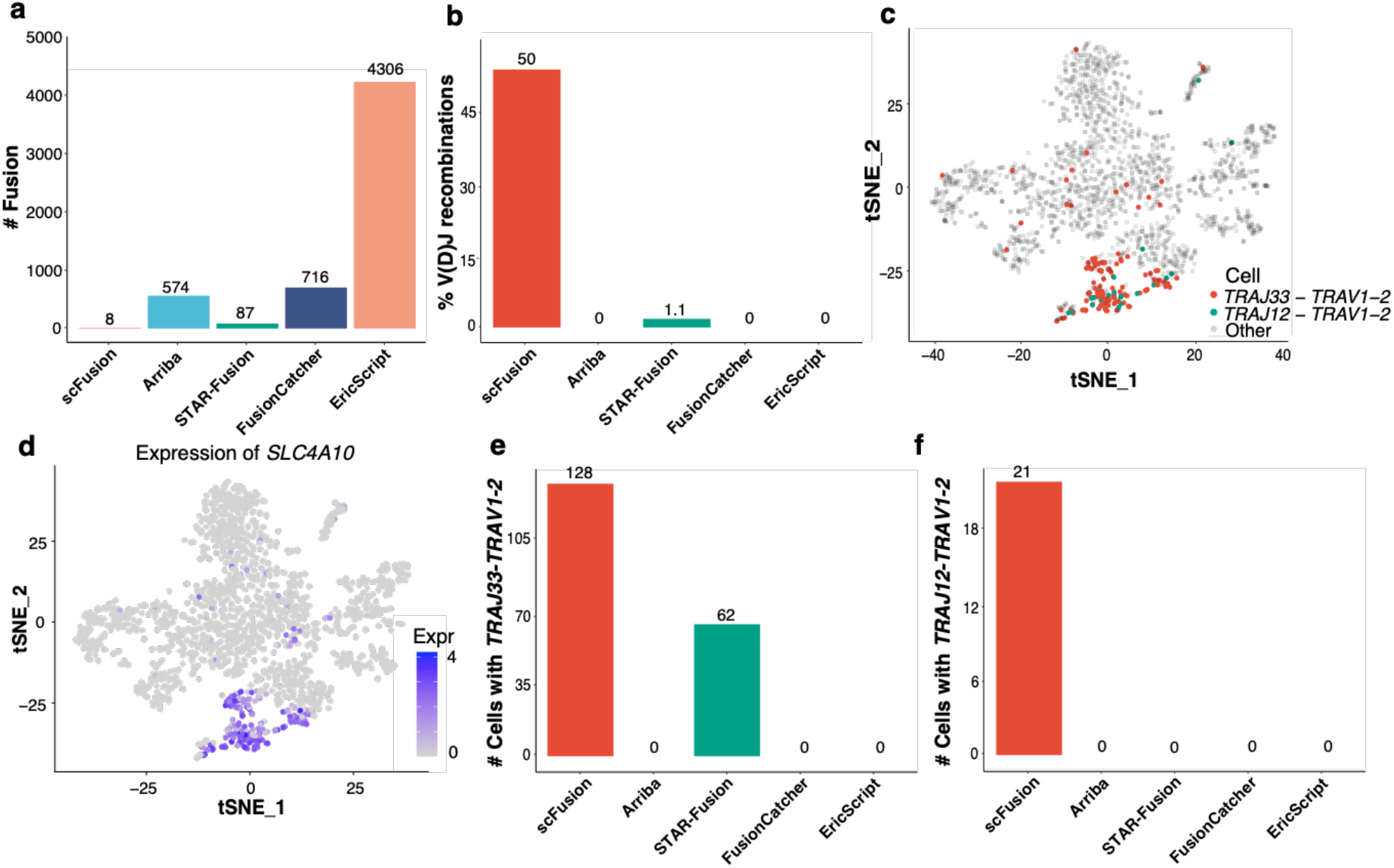
The T cell scRNA-seq data. (**a**) The number of detected gene fusions by the five methods. (**b**) The percentages of V(D)J recombinations in fusions detected by the five methods. (**c**) The expressions of *SLC4A10* shown in the tSNE plot of all T cells. (**d**) The cells with *TRAJ33-TRAV1-2* and *TRAJ12-TRAV1-2* colored in the tSNE plot. (**e**,**f**) The barplots of numbers of cells with the *TRAJ33-TRAV1-2* (e) and *TRAJ12-TRAV1-2* (f) recombinations by different algorithms.

### scFusion has superior computational efficiency in real scRNA-seq data

We first compared the computational time of different algorithms in real scRNA-seq data (Fig. 4). We considered three scRNA-seq data. The three scRNA-seq datasets are a newly sequenced spike-in scRNA-seq data (126 cells), a T cell data (2355 cells) and a multiple myeloma data (597 cells). Details about the data is described below. The computational time of scFusion is only about a half of Arriba and is only ∼10% or less of STAR-Fusion, EricScript and FusionCatcher, demonstrating superior computational efficiency of scFusion.

### scFusion detects fusions with less false discoveries in a single cell spike-in dataset

To test whether scFusion can detect fusions in real scRNA-seq data, we spiked-in 5 known gene fusions to single cells and performed single cell sequencing^8,9^ (**Methods** and Extended Data Fig. 6a). In total, we obtained scRNA-seq data of 126 single cells. scFusion reported 4 out of 5 spike-in gene fusions and 10 other fusions (Extended Data Table 1, 2). scFusion did not detect the spiked-in fusion *CCDC6-RET*, since the fusion occured in only two cells and did not achieve the significance level. The bulk methods also detected 4 or 5 spike-in fusions (Table S3-6), but they reported much more fusions (60, 2845, 353, and 898 for STAR-Fusion, EricScript, Arriba, and FusionCatcher, respectively), indicating potentially high FDRs. We also performed bulk RNA-seq for the spike-in data. 79% (11) of fusions detected by scFusion had 2 or more supporting chimeric reads in the bulk data, and only 9%-27% of fusions detected by the bulk methods had bulk supporting chimeric reads (Extended Data Fig. 6b), indicating that the FDR of scFusion is much lower.

**Table 1.** The contingency table between the *TRAJ33-TRAV1-2/TRAJ12-TRAV1-2* fusion and the expression of *SLC4A10*. The high expression are the cells whose *SLC4A10* expression is greater than 1 and the low expression are other cells. The Fisher’s exact test gives a p-value smaller than 10^−16^.

|  | *TRAJ33-TRAV1-2/TRAJ12-TRAV12* | Other |
| --- | --- | --- |
| High Expression | 108 | 90 |
| Low Expression | 40 | 2117 |

**Fig. 6.**
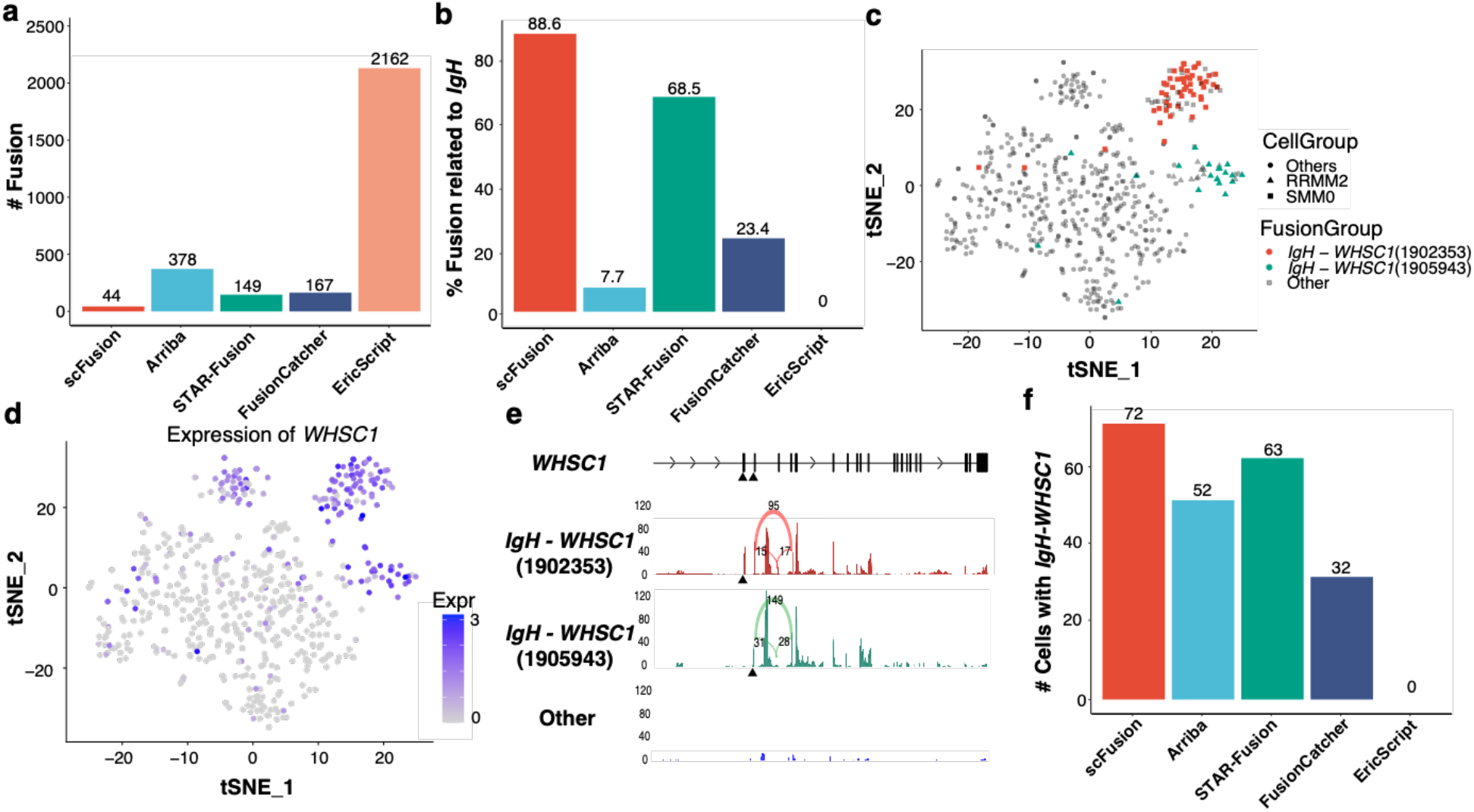
The MM scRNA-seq data. (**a**) The number of detected gene fusions by the five methods. (**b**)The percentage of IgH-related fusions in fusions detected by the five methods. (**c**) The tSNE plot the all MM single cells. The cells with two *IgH-WHSC1* fusions are colored in the plot. The cells from patient RMM2 and SMM0 are marked by triangle and rectangle, respectively. (**d**) The expression of *WHSC1* shown in the tSNE plot. (**e**) The mean read depth of *WHSC1* at different locations for the cells with the two *IgH-WHSC1* fusions and the cells without the fusions. The black triangles indicate the breakpoints of the two fusions. The supporting number of splicing junctions are also shown in the plot (the numbers above the arcs). The read depth of a single cell at a location is calculated as the number of reads covering the location per million. The mean read depth is the average depth of all cells in a group. (**f**) The barplots of numbers of cells with the *IgH-WHSC1* fusions by different algorithms.

### scFusion demonstrates high sensitivity in detecting marker fusions in a T cell dataset

The T cell data consists of 2,355 T cells from the immune data containing ∼7,000 immune cells profiled from patients with liver cancer ^22^. T cells of this data are non-malignant cells. Other than the V(D)J recombinations of TCR genes, gene fusions should be very rare in T cells. scFusion identified 8 gene fusions, much less than bulk methods (Fig. 5a, Extended Data Table 7-11). Among the fusions detected by scFusion, 6 involve TCR genes and 4 out of 8 (50%) are V(D)J recombinations, indicating that many of these candidates are likely true positives. In comparison, only 1.1% (1) of STAR-Fusion candidates are V(D)J recombinations and no candidates of other bulk methods are V(D)J recombinations (Fig. 5b). The two most frequent V(D)J recombinations given by scFusion are *TRAJ33-TRAV1-2* and *TRAJ12-TRAV1-2*, with 128 and 21 supporting cells, respectively. Mucosal-associated invariant T (MAIT) cells are known to express the invariant *TRAJ33-TRAV1-2* and *TRAJ12-TRAV1-2* TCR α-chain ^22,32^. Thus, the single cells with one of these fusions are likely MAIT cells. *SLC4A10*, a marker gene of MAIT cells ^33^, only expressed in a cluster of T cells and many of these T cells had the *TRAJ33-TRAV1-2* or *TRAJ12-TRAV1-2* fusion (Fig. 5c, d). Differential expression analysis between the cells with and without these recombinations identifies 70 upregulated genes in the cells with the recombinations, among which *TRAJ33, TRAV1-2* and *SLC4A10* are the top three most significantly upregulated genes (Extended Data Fig. 7a, b). Further, the *SLC4A10* expression are highly significantly associated with the fusion (Fisher’s exact test, P-value < 10^−16^; Table 1). These results indicate that the *TRAJ33-TRAV1-2* and *TRAJ12-TRAV1-2* fusions identified by scFusion are *bona fide* fusions and the cells with either fusion are MAIT cells. MAIT cells are an important component of the immune system. However, until recently, the detection of MAIT cells still relies on the combinations of cell markers and it is unclear how well these markers define the MAIT cells in different tissues or diseases^34^. scFusion provides an alternative way to sensitively define the MAIT cells. STAR-Fusion was the only bulk method that identified the *TRAJ33-TRAV1-2* fusion (62 cells) (Fig. 5e), and no bulk method reported any cells having the *TRAJ12-TRAV1-2* fusion (Fig. 5f), indicating that scFusion is more powerful in detecting shared fusions among single cells.

### Senstitive fusion detection by scFusion provides mechanistic insights in a multiple myeloma dataset

Finally, we applied scFusion to a multiple myeloma (MM) dataset consisting of 597 MM single cells from 15 patients ^28^. MM is a cancer of plasma cells. Approximately half of the myelomas have immunoglobulin heavy (*IgH*) chain translocations and around 10% of the myelomas have translocations involving immunoglobulin lambda (*IgL*) light chain locus ^35,36^. In this dataset, scFusion identified 44 fusions and the bulk methods identified much more fusions (Fig. 6a, Extended Data Table 12-16). Around 88.6% (39) of scFusion candidates involve immunoglobulin genes (including 33 recombinations of immunoglobin genes), much higher than that of bulk methods. (Extended Data Table 12, Fig. 6b). scFusion successfully identified the recurrent *IgH-WHSC1* fusion in MM with two different breakpoints at *WHSC1*, 1902353 and 1905943 at chromosome 4 (Extended Data Table 12). The *IgH-WHSC1* fusions are in-frame fusions of *WHSC1*. All 52 cells with the breakpoint at 1902353 are from the patient SMM0. Similarly, all 20 cells with the other breakpoint are from the patient RRMM2 (Fig. 6c). The expression of *WHSC1* is significantly higher in cells with the *WHSC1* fusions than other cells (Fig. 6d). Interestingly, in cells with the *WHSC1* fusions, the sequencing coverage of *WHSC1* sharply increases at the downstream of the breakpoints, but the sequencing coverage of *WHSC1* in other cells largely keeps constant, indicating that the fusions probably lead to the over-expression of *WHSC1* (Fig. 6e, Extended Data Fig. 7c). *WHSC1*, also known as *NSD2*, is one of the most commonly fused genes with IgH in multiple myeloma^37-39^. *WHSC1* is a known oncogene listed in COSMIC ^40^. Overexpression of *WHSC1* drives the chromatin change in an H3K36me2-dependent manner^41^. Differential expression analysis found 115 upregulated genes and 12 down-regulated genes in the cells with the *IgH-WHSC1* fusions (q-value < 0.05 and log2 fold change < -0.5 or > 0.5) (Extended Data Fig. 7d, Extended Data Table 17). The upregulated genes include genes known be co-expressed with *WHSC1* such as *MAL* ^42^ and *SCARNA22* ^43^. The downregulated genes include known oncogenes in MM such as *CCND1* and *FRZB*. In fact, *CCND1* and *FRZB* tend to only express in cells without the *IgH-WHSC1* fusions (Extended Data Fig. 7e, f). The bulk methods also detected the *IgH-WHSC1* fusions, but they reported less number of cells with the fusions than scFusion (Fig. 6f) and potentially much more false positives (Fig. 6a, b). The large number of false positives given by the bulk methods makes it very difficult for downstream analysis, thus limiting their application in single cell analyses. In comparion, scFusion has much fewer false discoveries and is more powerful in detecting important shared fusions such as *IgH-WHSC1*.

## Discussion

We describe a single cell gene fusion detection method scFusion by integrating fusion signals from multiple cells. The major challenge of fusion detection in single cells is the large number of false positives. To address this challenge, we introduced a statistical testing procedure to control the FDR. This procedure assumes most of fusion signals are technical noises and true fusions generally have stronger signal than technical noises. In addition, some systematic technical artefacts can occur recurrently in multiple single cells and might be assigned with a significant p-value by the statistical procedure. We developed a deep learning model to learn the unknown technical artefacts and to filter the false postives generated by these artefacts.

We evaluated the performance of scFusion using simulation data and three real scRNA-seq datasets. In the simulation, a critical problem is to generate technical chimeric artefacts. Unfortunately, the mechanisms generating these artefacts are largely unknown and it is difficult to generate chimeric artefacts that have the same pattern as those in the real scRNA-seq data. We bypassed this problem by taking chimeric reads from real scRNA-seq data and adding them to the simulation data. In this way, we guaranteed that the simulation faithfully reflects the performance of fusion detection methods in real data. scFusion outperformed the bulk methods in terms of precision and F-score. In the real data analysis, we found that scFusion generally detected much less fusions compared with the bulk methods. Especially, scFusion only detected 8 fusions in the T cell data while the bulk methods detected much more fusions. Considering that the T cells are normal cells, they should not have many fusions, implying that many fusions detected by the bulk methods are false positives. In addition, among the fusions reported by the bulk methods, many were very likely to be technical artefacts (Extended Data Fig. 8). For example, the *GADD45G--HSPH1* fusion was reported by all four bulk methods, but it contained a poly-A subsequence and its artefact score was almost 1.

We demonstrated application of scFusion in two different scRNA-seq datasets. In both datasets, scFusion was able to identify subgroups of single cells with specific gene fusions, i.e. the MAIT cells with the invariant TCR recombinations and the MM cells with the *IgH-WHSC1* fusions. MAIT cells in human are known to express three TCR α-chains, of which the most abundant is *TRAJ33-TRAV1-2* ^34^. scFusion identified two of the three and found that *TRAJ33-TRAV1-2* about 5 times more abundant than *TRAJ12-TRAV1-2*. The *TRAJ20-TRAV1-2* was not identified possibly because it was rare in MAIT cells. Gene differential expression analysis discovered 15 genes significantly differentially expressed between *TRAJ33-TRAV1-2* and *TRAJ12-TRAV1-2* cells (Table S18). As expected, the expressions of *TRAJ33* and *TRAJ12* are higher in *TRAJ33-TRAV1-2* and *TRAJ12-TRAV1-2* cells, respectively. Interestingly, *ALPK1* and *TIFA* are highly expressed in the cells with *TRAJ12-TRAV1-2* (Extended Data Fig. 9). Recent studies^44^ show that *ALPK1-TIFA* axis is a core innate immune pathway against patheogens such as *Helicobactor pylori*, implying that MAIT cells expressing different TCR α-chains might have different roles in the immune system. In the MM data, the *IgH-WHSC1* fusions are the most significant immunoglobulin-related fusions reported by scFusion, leading to the overexpression of *WHSC1. WHSC1* plays an important role in cell development. Overactivating *WHSC1* promotes the uncontrolled growth and division of cancer cells ^45^. In the MM data, *WHSC1* is highly expressed in single cells from three patients (SMM0, RRMM1 and RRMM2). scFusion did not find *WHSC1* fusions in single cells from RRMM2. It is likely that the expression of WHSC1 was activated in RRMM2 by mechnisms other than *WHSC1* fusions or that the fusion was missed by scFusion. Taken

together, these applications show that accurate gene fusion detection at single cell level could provide key information for defining cellular subtypes or subclones, which might have important implications for many applications, such as identifying drug resistant or sensitive subclones in tumor.

Gene fusions are one of the most widely studied biological events in cancer. Analysis of gene fusions at single cell level will provide unprecedent opportunities to study their roles in tumor development, tumor heterogeneity as well as tumor cell’s responses to various pharmatheutical therapies. We expect that scFusion will find its application in many different single cell researches.

## Methods

### Statistical Model

Let *y*_*ij*_ be the number of split-mapped reads supporting a fusion candidate *i* in a cell *j*. Denote *Y*_*ij*_ the random variable corresponding to *y*_*ij*_. Theoretically, the random variable *Y*_*ij*_ should follow a mixture distribution. One component of the mixture distribution is the distribution of supporting reads of true gene fusions and another is the distribution of supporting reads of the technical chimeric artefacts. However, since the candidate fusions have very few true fusions, we can safely use all *y*_*ij*_ to estimate the distribution of split-mapped reads from technical chimeric artefacts. With the estimation of this background distribution, we can then perform statistical tests to identify candidates that are unlikely to be generated from the background distribution. We assume that *Y*_*ij*_ follows a zero-inflated negative binomial (ZINB) distribution^46^. The ZINB distribution has three parameters. One parameter is *p*_*i*_, the probability that the cell *j* has no split-mapped read supporting the fusion candidate *i* or *p*_*i*_ = *P*(*Y*_*ij*_ = 0). The other two parameters of the ZINB are the mean *μ*_*ij*_ and the overdispersion parameter *λ* of the negative binomial (NB) distribution. Thus, we have *Y*_*ij*_∼*ZINB*(*p*_*i*_, *μ*_*ij*_, *λ*). Conditional on *Y*_*ij*_ > 0, *Y*_*ij*_ follows a zero-truncated NB distribution. The ZINB distribution could depend on other factors, such as the expression levels of the partner genes and the GC content ^47^. Analysis of several different scRNA-seq data shows that the number of technical chimeric artefacts indeed strongly depends on the gene expression and the GC content (Fig. 2a, b). Let *EL*_*ij*_ < *EH*_*ij*_ be the expressions of the two partner genes corresponding to *Y*_*ij*_ and *GC*_*i*_ be the GC content of the exonic sequence near the junction of the candidate gene (200bp). Further, define 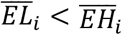 as the mean expression of the two partner genes of the candidate fusion *i* across all single cells. We consider the following model

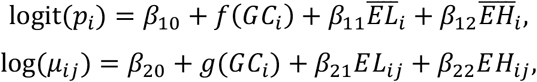

where *f* and *g* are unknown functions, and the ***β***_1_ = (*β*_10_, *β*_11_, *β*_12_) and ***β***_2_ = (*β*_20_, *β*_21_, *β*_22_) are unknown parameters. We represent *f* and *g* using spline functions, 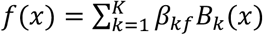 and 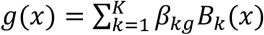, where *K* is the number of spline base functions (by default K is set as 5). Denote ***θ*** = (***β***_1_, ***β***_2_, ***β***_*f*_, ***β***_*g*_) with ***β***_*f*_ = (*β*_1*f*_, ⋯ *β*_*kf*_) and ***β***_*g*_ = (*β*_1*g*_, ⋯ *β*_*kg*_). We estimate the unknown parameters ***θ*** by maximizing the following likelihood,

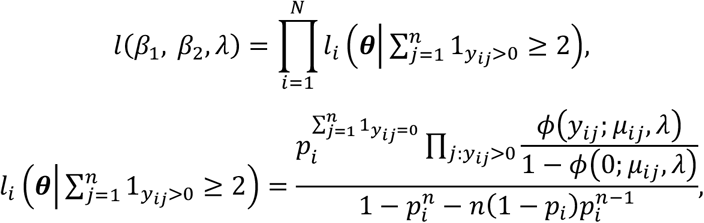

where *N* is the number of fusion candidates supported by at least 2 cells, *n* is the number of cells, *ϕ* is the probability density function of NB distribution with mean *μ*_*ij*_ and overdispersion *λ*. Note that to reduce computational burden, we only consider the candidates with at least two cells having supporting split-mapped reads and thus the likelihood *l*_*i*_ for the candidate *i* is a conditional probability.

### Statistical Test for Significant Fusions

After estimating the unknown functions and unknown parameters, we have the parameter estimates 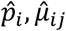 and 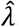. We use these estimates and a resampling scheme to obtain a p-value for each fusion candidate. Specifically, for each fusion candidate *i*, we sample 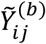 from 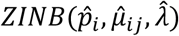 for each cell *j* and obtain the sum 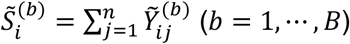. Denote 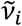 and 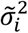 be the sample mean and variance of 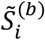, respectively. The distribution of 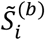 can be approximated by a normal distribution with the mean and variance approximately 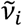 and 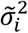, respectively. Let 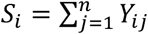. The p-value for the fusion candidate *i* is set as 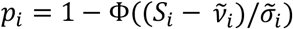, where Ф is the cumulative distribution of the standard normal distribution. We set *B* = 1,000 in all analyses. To determine a p-value cutoff to control the FDR, we split our candidate fusion set to two subsets. The first subset is a high-quality subset which more likely contains the true gene fusions and the second subset is the remaining candidates. Suppose that *n*_1_ and *n*_2_ as the total number of candidates in the first and the second subset, respectively. Given a p-value cut-off *c*, let *m*_1_(*c*) and *m*_2_(c) be the number of candidates in the two subsets with p-values less *c*, respectively. Since the second subset contains much fewer true positives than the first subset, 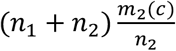 can be used as an estimate of total number of false discoveries at the p-value cutoff *c* and thus the FDR is roughly 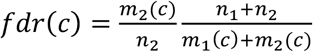 We choose *c* such that *fdr*(*c*) ≤ *α* for a given *α* (usually 0.05). In generally, we believe that candidates with more supporting cells are more likely to be true. Hence, we choose the first subset as the fusion candidates with at least 1% of the total cells having the fusion and with at least *s* supporting reads per cell on average, where *s* is taken as 1.25 in all simulation and real data analysis.

### The bi-LSTM

Each single cell usually has over 10,000 chimeric reads (Extended Data Fig. 3c) and the vast majority of them are obviously not signals of gene fusions but technical artefacts. We set the negative training data as the subsequences from chimeric reads, representing the technical artefacts. An ideal positive training data would be chimeric reads from true fusions. Unfortunately, we do not know which chimeric reads are from true fusions. Even if we knew some, the number of such reads would be too small to train the bi-LSTM network. To overcome this challenge, a proxy task strategy is applied: we set the positive training data as sequences generated by concatenating random pairs of short reads. Since the sequences are randomly generated, they should not contain features of technical chimeric reads and hence characteristics of technical artefacts could be learned by comparing with these random sequences. More specifically, the negative training data of the bi-LSTM is chosen as the subsequences (60 bp) of chimeric reads covering the junction positions and the junction positions are required to be at 15-45 bp of the subsequences. The positive training data is the similar 60bp subsequences of randomly concatenated short read pairs.

In the bi-LSTM (Extended Data Fig. 2b), the DNA sequences, with 60 nucleotides and one fusion site, are given as the input to the embedding layer. The four types of nucleotide (i.e., A, T, G and C) and the fusion site are represented by five different 5-dimensional feature vectors. The output of the embedding layer is passed on to three sequence-to-sequence bi-LSTM layers (with 32, 64, and 128 bi-LSTM units, respectively) and further to a sequence-to-one bi-LSTM layer for sequential feature extraction. The extracted features are fed to two fully connected layers and finally to a softmax layer to produce the softmax probabilities of the read being classified to technical chimeric artefacts. In the training process, the binary cross-entropy is used as the loss function and model parameters are updated using the Adam optimizer^48^. The model is pre-trained for 200 epochs with batch size set to 500. The retraining is trained for 30 epochs with the pre-trained model as initial value.

Using this bi-LSTM model, scFusion gives each fusion candidate a technical artefact score. By default, fusions with scores greater than 0.75 are filtered. The training step is computationally expensive. To expediate the training, scFusion provides a pre-trained model that can be used as the initial value to train bi-LSTM models for new datasets. Note that if we directly used the pre-trained model, the median AUC and AUPR were 0.673 and 0.749 (Extended Data Fig. 2h, i), respectively. The convolutional neural network (CNN) is another popular neural network model used in many biological applications^49^. We also built a CNN model for comparison and found that the bi-LSTM generally performed better than the CNN (Supplementary Material, Extended Data Fig. 10). Hence, we use the bi-LSTM in all data analyses.

### Simulation setup

The simulation data was generated using FusionSimulatorToolkit ^19^. This toolkit can simulate RNA-seq data with gene fusions by learning from a real RNA-Seq dataset such as its expression, insert size, read length and mutation rate. We provided the immune cell scRNA-seq^22^ data to FusionSimulatorToolkit to generate RNA-seq data with 150 simulated fusions at various gene expression levels. However, the patterns of technical chimeric reads are unknown. FusionSimulatorToolkit and other available RNA-seq data simulators cannot generate chimeric reads mimicking those in real scRNA-seq data. Observing that immune cells are normal cells, other than chimeric reads generated from recombinations of T Cell Receptor (TCR) genes in T cells and immunoglobulin genes in B cells, almost all other chimeric reads should come from technical artefacts. Thus, we randomly chose chimeric reads not involving TCR and immunoglobulin genes from the immune cell scRNA-seq data and added them to the simulated data. Chimeric reads from one cell’s data were only added to simulation data of one cell. This simulation process guaranteed that the chimeric reads in simulation data were similar to those in real scRNA-seq data and that the simulation faithfully reflected the performance of scFusion in real scRNA-seq data.

### Gene fusion spike-in and single cell sequencing

Five fusion genes cDNA sequences were synthesized and constructed into lentiviral vectors respectively. To construct a cell line with stable expression of fusion genes, every lentiviral vector and two auxiliary packaging plasmids were co-transfected into the 293T cells. After 48 hours, the supernatant was collected from the 293T cells and filtered through a 0.45 uM membrane. The five different recombinant lentiviral particles containing the target fusion genes and the green fluorescent protein (GFP) reporter gene were collected. Then, the five recombinant lentivirus particles infected into 293T cells, respectively. 72 hours after infection, the medium was changed and the expression of GFP in cells was checked under a fluorescence microscope for determining if lentivirus infection was successful. After the infection, five cell lines expressing the target fusion genes were collected. The single GFP positive cells were filtered into a 96-well PCR plate by fluorescence activated cell sorting (FACS) for the single-cell RNA library construction. Poly(A)-transcripts of total RNA of single cells were reverse transcribed and amplified using the SMART-seq2 ^8,9^ protocol. The amplified cDNA was tagmented by Nextera XT kit (Illumina) and libraries were sequenced by NovaSeq (Illumina). The 293T cells were purchased from the National Infrastructure of Cell Line Resource (http://www.cellresource.cn/).

### scRNA-seq data analysis

For scFusion, short reads are first aligned with STAR(v 2.7.4a) to the human reference genome (hg19). scFusion only considers short reads mapped to uniquely mappable positions in the exonic regions of the reference genome (uniquely mappable for 75bp sequences). Then, the split-mapped reads are clustered. If the breakpoints of two split-mapped reads are no larger than 3 bp away from each other, the two split-mapped reads are clustered together. A fusion candidate’s breakpoint is taken as the median of the breakpoint positions of its all supporing chimeric reads.

Arriba(v1.0.1), EricScript(v0.5.5b), STAR-Fusion(v1.8.1), FusionCatcher(v1.10) were all runned on the default parameters. The reference genome was always chosen as hg19. For the bulk methods, we also applied the same *ad hoc* filters of scFusion to make different algorithms comparable. Specifically, we applied the pseudogene-, lncRNA-, no-approved-symbol-gene-, intron-, too-many-partner- and too-many-discordant-filters to the fusion candidates of bulk methods.

### Gene expression analysis

The expression matrix of T cell and multiple myeloma dataset were downloaded from the Gene Expression Omnibus (GEO) (https://www.ncbi.nlm.nih.gov/geo/). First, we used Seurat (v 3.2.0) ^50,51^ to read the Transcripts Per Million (TPM) matrix. The expression was further normalized using the NormalizeData function in Seurat. The highly variable genes were identified using the function FindVariableGenes. Their expression was scaled and centered along each gene using ScaleData. Then we performed dimension reduction using principal component (PC) analysis. We selected the first 30 PCs for t-distributed stochastic neighbor embedding (tSNE), and tSNE plots were generated using Seurat. To identify differentially expressed genes, we used the function FindAllMarkers in Seurat with the Wilcoxon rank sum test. Genes expressed in at least 10% cells within the cluster and with the log fold change more than 0.5 and the adjusted p-value smaller than 0.05 were considered as differentially expressed genes.

## Data availability

Raw data of the spike-in data will be available at the Genome Sequence Archive (GSA) of BIG Data Center (http://bigd.big.ac.cn/gsa), Beijing Institute of Genomics (BIG), Chinese Academy of Sciences, with an accession number HRA000343. The scRNA-seq data GSE81812, GSE99795, GSE110499, GSE118900, GSE127298, GSE140228 were obtained from the Gene Expression Omnibus (GEO) (https://www.ncbi.nlm.nih.gov/geo/).

## Code availability

scFusion is available at github (https://github.com/XiDsLab/scFusion).

## Acknowledgements

This work was supported by the National Natural Science Foundation of China [11971039, 11471022 and 71532001] and the Recruitment Program of Global Youth Experts of China.

## Author contributions

R.X. conceived and supervised the study. Z.J. developed the statistical model. W.H., Z.J. and X.W. developed the deep-learning model. J.L. performed the the single cell experiment. Z.J., R.X. and N.S. performed data analysis. R.X., Z.J., N.S., W.H. and X.W. wrote the manuscript, with feedback from P.P.

