## Supplemental text and figures for "Single cell gene fusion detection by scFusion"

**This PDF file includes:**

Supplementary text
Extended Data Figures 1 to 10
SI References

**Other supplementary materials for this manuscript include the following:**

Extended Data Table 1 to 18

### Supplementary Information Text

#### The convolutional neural network model

The convolutional neural network (CNN) model is mainly composed of three layers: a convolution layer, a pooling layer and a fully connected layer (Figure S4A). The input of the network is a 60bp read, which contains the bases near the breakpoint of the fusion gene, and an integer between 15 and 45 that indicates the position of the breakpoint. The output is a probability that the sequence is a technical chimeric artefact.

We first convert the input sequence to a padded 5-row array that each row represents one of four bases (A, G, T, C) or the breakpoint position. Specifically, given the size of convolution kernel m, we convert a sequence $S$ of length n to a 5*(n+2m-2) array $A$ in this way:

$$A_{i,j} = \begin{cases} 0.25 & if \ \ i = 1...4 \ and \ j < m \ or \ j > n - m \\ 1 & if \ (i, S_{j-m+1}) \ = (1, A) \ or \ (2, G) \ or \ (3, C) \ or \ (4, T) \\ 1 & if \ i = 5 \ and \ the \ breakpoint \ position \ is \ j - m + 1 \\ 0 & otherwise \end{cases}$$

where $A_{i,j}$ denotes the (i, j) element of $A$ and $S_i$ denotes the i-th base of $S$.

The convolutional layer takes the array as input, and each convolution kernel performs one-dimensional convolution on this 5-channel input. The number of kernels as well as its size are tunable parameters. In our model, we use 2048 convolution kernels of length 16 ($m = 16$) by default. Therefore, the output of this layer is of size 2048*75. The output feature map of each kernel is then rectified by adding a bias and being activated by ReLU.

The pooling layer performs max operation on the rectified features, aiming to integrate the features obtained by the convolution. After this stage, we get a 2048-dimensional vector. Finally, the feature vector is passed to the fully connected layer, which consists of a hidden layer and an output layer. The hidden layer is of 256 neurons by default and the output layer is a single neuron with Sigmoid function.

We use negative log-likelihood as the loss function, and Adam[1] with L2 regularization for optimization. We use mini-batch to speed up training and add a dropout layer after convolution to reduce overfitting. By default, batch size, learning rate, weight decay and dropout rate is set to 512, $3*10^{-3}$, $1*10^{-4}$ and 0.25 respectively.

**Commands**

We list how we use the bulk method in the following.

**STAR mapping**:

*STAR --runThreadN 20 --genomeDir CTAT_GENOME_LIB/GRCh37_ gencode_v19_CTAT_lib_Feb092018/ctat_genome_lib_ build_dir/ref_genome.fa.star.idx --readFilesIn /1_1.fastq 1_2.fastq --outSAMtype BAM SortedByCoordinate --chimOutType SeparateSAMold --outSAMunmapped Within KeepPairs --quantMode GeneCounts --outFileNamePrefix 1/human --chimSegmentMin 12 --chimJunctionOverhangMin 12 --alignSJDBoverhangMin 10 --alignMatesGapMax 100000 --alignIntronMax 100000 --chimSegmentReadGapMax 3 --alignSJstitchMismatchNmax 5 -1 5 5*

**STAR-Fusion:**

*STAR-Fusion-v1.8.1/STAR-Fusion --left_fq 1_1.fastq --right_fq 1_2.fastq --genome_lib_dir /GRCh37_gencode_v19_CTAT_lib_Oct012019.plug-n-play/ctat_genome_lib_build_dir/ --CPU 20 --output_dir STAR_Fusion_Res/*

**EricScript:**

*Ericscript/ericscript.pl -db Ericscript/ericscript_db_homosapiens_ensembl73 -p 20 -name 1 -o EricRes/1 1_1.fastq 1_2.fastq*

**Arriba:**

Arriba is piped with STAR:

*STAR --runThreadN 20 --genomeDir CTAT_GENOME_LIB/GRCh37_ gencode_v19_CTAT_lib_Feb092018/ctat_genome_lib_ build_dir/ref_genome.fa.star.idx --genomeLoad NoSharedMemory --readFilesIn 1_1.fastq 1_2.fastq --outStd BAM_Unsorted --outFileNamePrefix OUTDIR --outSAMtype BAM Unsorted --outSAMunmapped Within --outBAMcompression 0 --outFilterMultimapNmax 1 --outFilterMismatchNmax 3 --chimSegmentMin 10 --chimOutType WithinBAM SoftClip --chimJunctionOverhangMin 10 --chimScoreMin 1 --chimScoreDropMax 30 --chimScoreJunctionNonGTAG 0 --chimScoreSeparation 1 --alignSJstitchMismatchNmax 5 -1 5 5 --chimSegmentReadGapMax 3 | arriba -x /dev/stdin -o OUTDIR/fusions.tsv -O OUTDIR/fusions.discarded.tsv -a "$ASSEMBLY_FA" -g "$ANNOTATION_GTF" -b "$BLACKLIST_TSV" -T -P*

**FusionCatcher:**

*FusionCatcher/bin/fusioncatcher -d FusionCatcher/data/human_v95 -i 1 1.fastq,1_2.fastq -o FusionCatcherRes*

### Figures

**Extended Data Fig. 1:** Number of fusion candidates after applying different filters.

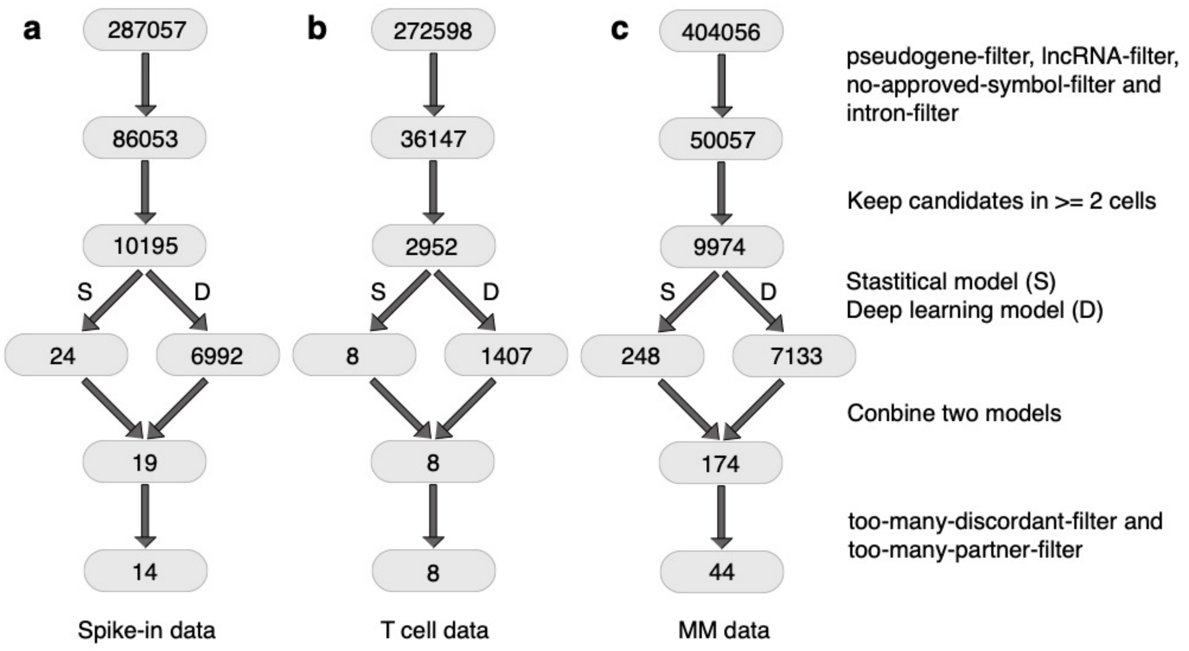


Number of fusion candidates after applying different filters for the (**a**) spike-in data, (**b**) T cell data and (**c**) MM data. The top is the total number of fusion candidates, and the second layer is the number of fusion candidates excluding pseudogenes, lncRNAs, genes without an approved symbol, and candidates in intronic regions (they are called pseudogene-, lncRNA-, no-approved-symbol- and intron- filters). The third layer is the number of candidates supported by at least 2 cells. The fourth layer shows the numbers of fusion candidates remained after the filtering by the statistical model and the deep learning model, respectively. The fifth layer is the number of candidates that pass both models. The last number indicates the number of output fusions after removing candidates whose number of supporting discordant reads is ten times more than supporting split-mapped reads (too-many-discordant-filter) and candidates with a partner gene involving in more than five fusion candidates (too-many-partner-filter).

**Extended Data Fig. 2:** The bi-LSTM model.

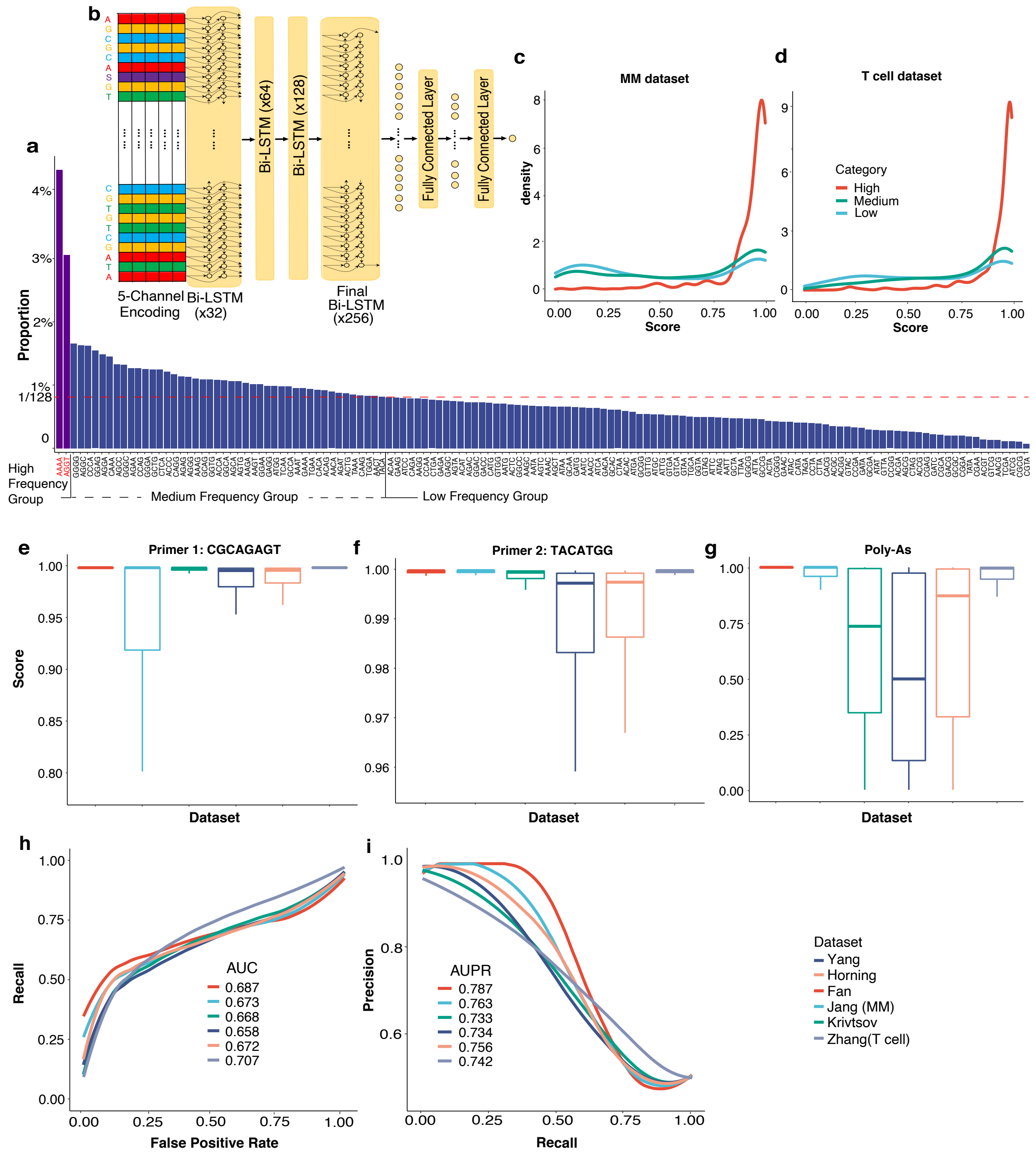


(**a**) The occurrence frequencies of the 128 possible 4-bp sub-sequences around the junction of chimeric reads. The red dashed line is the expected occurrence frequency if the 4bp sequence occurred randomly. (**b**) The design of the bi-LSTM. (**c,d**) The distributions of artefact scores of chimeric reads in the high, middle and low frequency group. The high frequency group consists of chimeric reads with AAAA and AGGT as their junction sequences (frequency > 3%). The low frequency group consists of chimeric reads whose 4bp junction sequences occurred in less than 1/128 of all chimeric reads. The medium frequency group is the other chimeric reads. (**c**) The MM dataset. (**d**) The T cell data. (**e,f,g**) The boxplots of artefact scores of reads containing (**e**) CGCAGAGT, (**f**) TACATGG, and (**g**) poly-As in six datasets. The sequences CGCAGAGT and TACATGG are often used as primer[2,3] sequences. These chimeric reads might be generated by random annealing of complementary sequences. (**h**)The ROCs and (**i**) PR curves of the Bi-LSTM without retraining.

**Extended Data Fig. 3:** Statistics of chimeric reads.

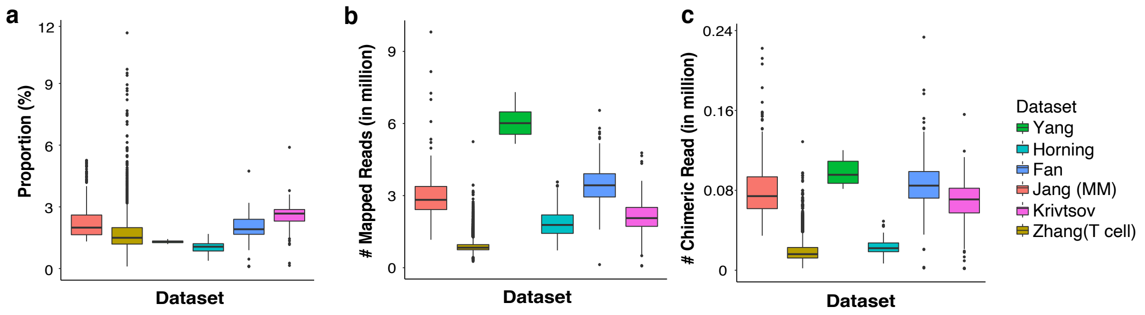


The boxplots of (**a**) the proportions of chimeric reads in all mapped reads, (**b**) the total number of mapped reads in different scRNA-seq data and (**c**) the total number of chimeric reads.

**Extended Data Fig. 4:** The precisions and recalls of scFusion and four bulk methods in six different simulation setups.

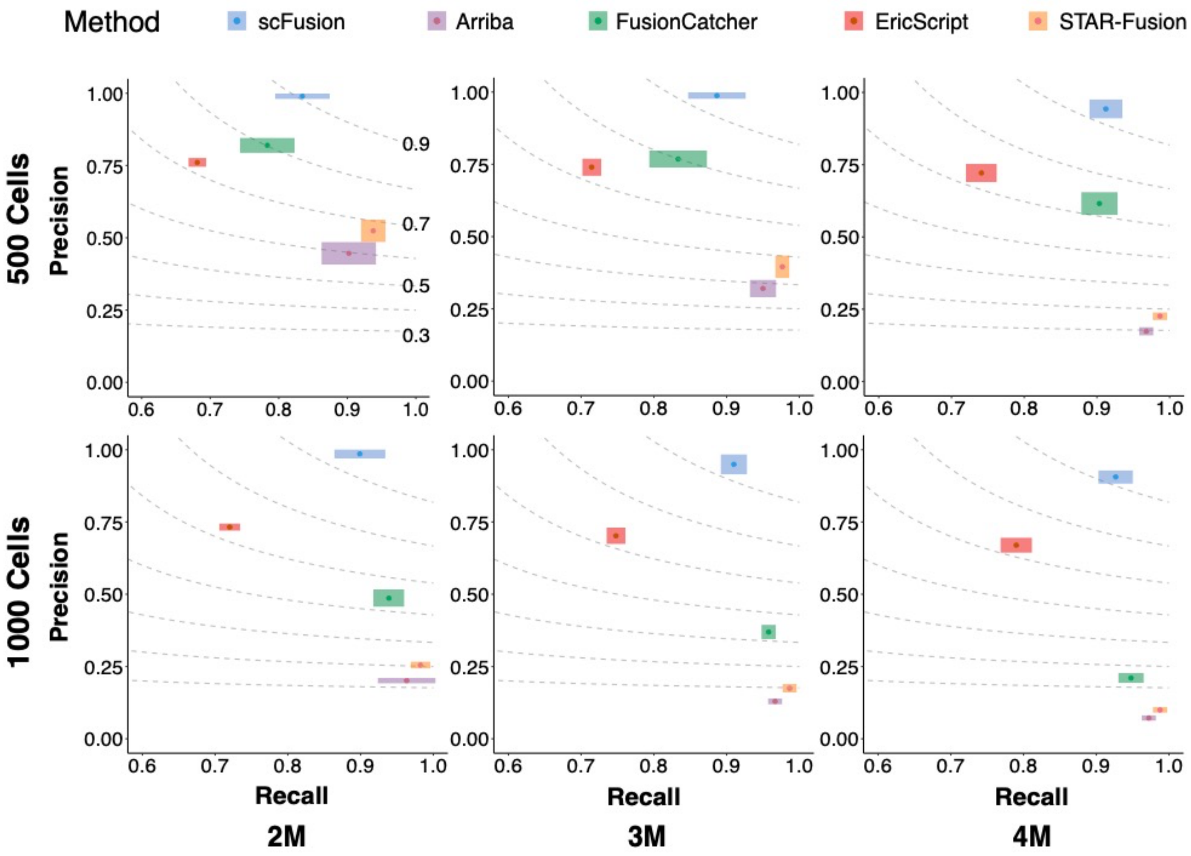


The figures in the two rows correspond to simulations with 500 cells and 1000 cells, and the figures in the three columns correspond to simulations with 2 million, 3 million and 4 million reads in each data. The dots in the figures are the means of precisions and recalls of ten simulations in each setup and the boxes ±1 standard deviation (SD) of the precisions and recalls. The dashed lines are the contour lines with constant F-scores (F-scores are marked in the top-left figure).

**Extended Data Fig. 5:** The detection powers of fusions at different expression levels

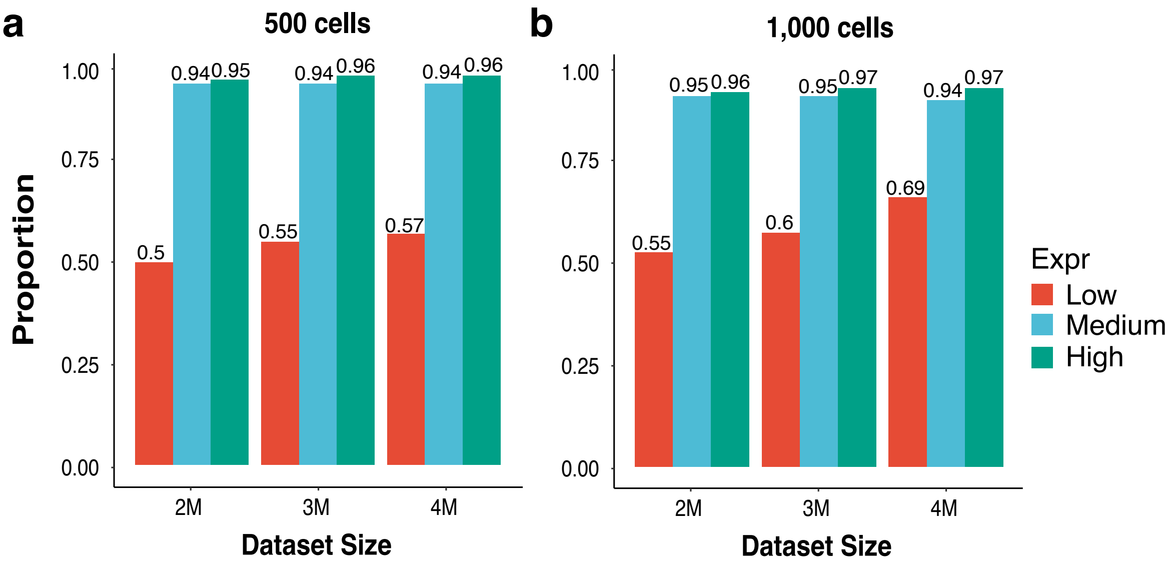


The detection powers of fusions at different expression levels using (**a**) 500 and (**b**) 1000 cells.

**Extended Data Fig. 6:** The spike-in data.

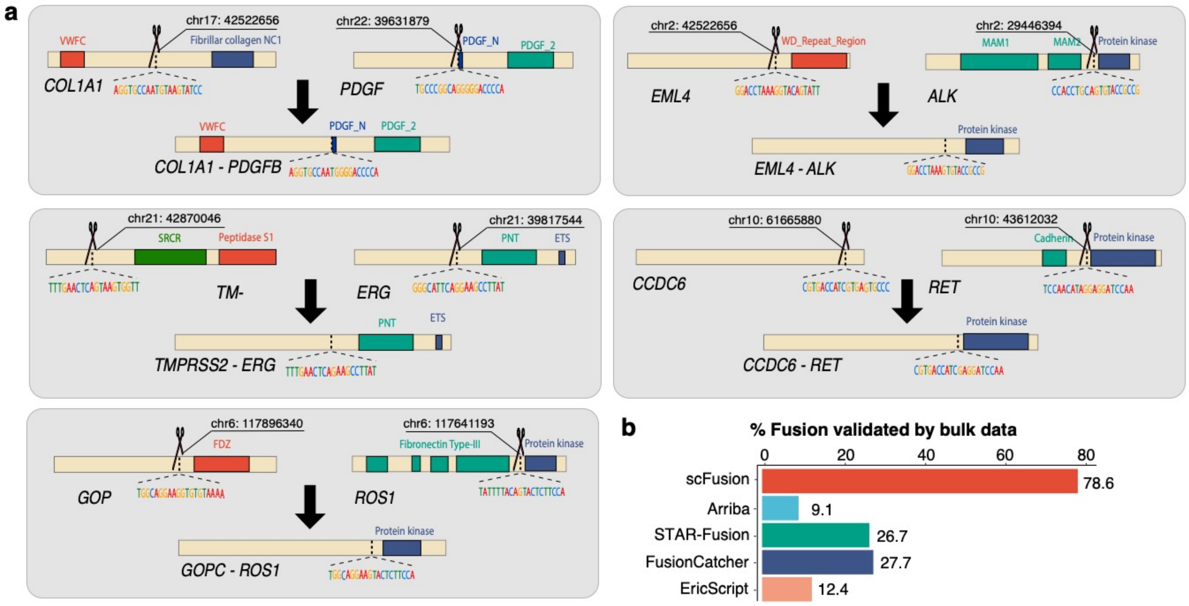


(**a**) The design of the five spike-in fusion genes. In each panel, the original partner genes are shown at the top and the fusion genes are shown at the bottom. The scissors represent the breakpoints of the fusions and the local sequences are shown below the genes and the fusion genes. (**b**) The percentage of reported gene fusions having supporting reads in bulk data.

**Extended Data Fig. 7:** Fusions are correlated with gene up-regulations.

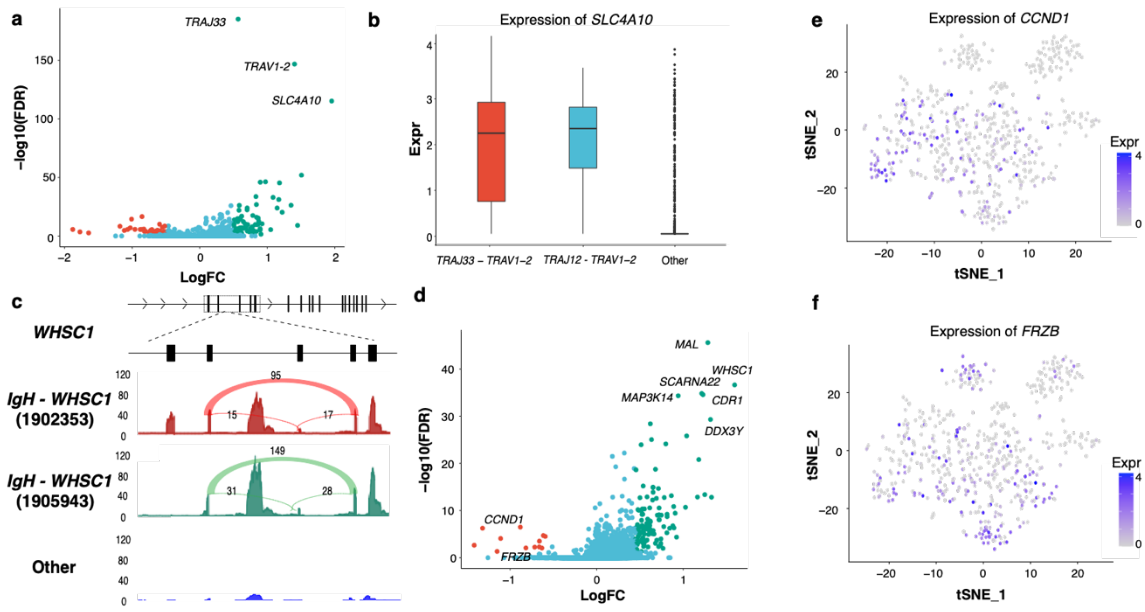


(**a**) The volcano plots of the differential expression analysis of T cells between cells with and without the *TRAJ33-TRAV1-2* or *TRAJ12-TRAV1-2* recombination. (**b**) The *SLC4A10* expression is significantly higher in cells with the *TRAJ33-TRAV1-2* or *TRAJ12-TRAV1*-2 recombination than other cells. (**c**) The local view of the mean read depth of *WHSC1* at different locations for the cells with the two *IgH-WHSC1* fusions and the cells without the fusions. (**d**) The volcano plots of the differential expression analysis of T cells between cells with and without the *IgH-WHSC1* fusion. (**e**) The expression of *CCND1* shown in the tSNE plot. (**f**) The expression of *FRZB* shown in the tSNE plot.

**Extended Data Fig. 8:** Six examples of gene fusions with high artefact scores.

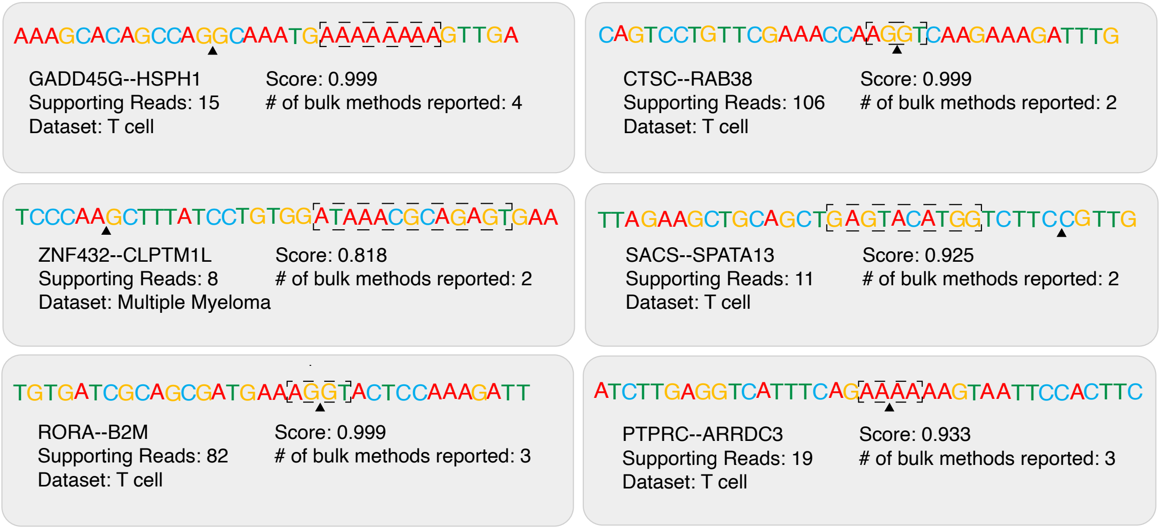


The black triangles show the junctions of gene fusions. The patterns in the dashed boxes might explain their high artefact scores.

**Extended Data Fig. 9:** Differential expression analysis.

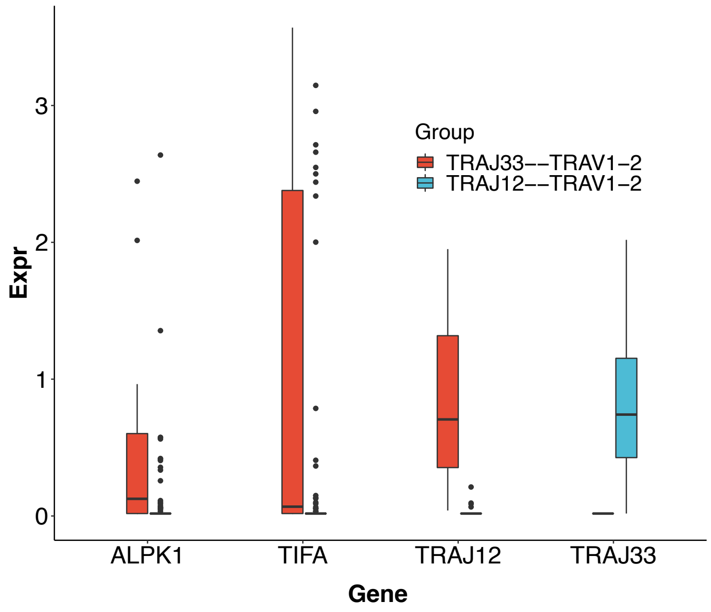


The expression of *ALPK1*, *TIFA, TRAJ33* and *TRAJ12* in the two cell groups.

**Extended Data Fig. 10:** Performance of CNN model.

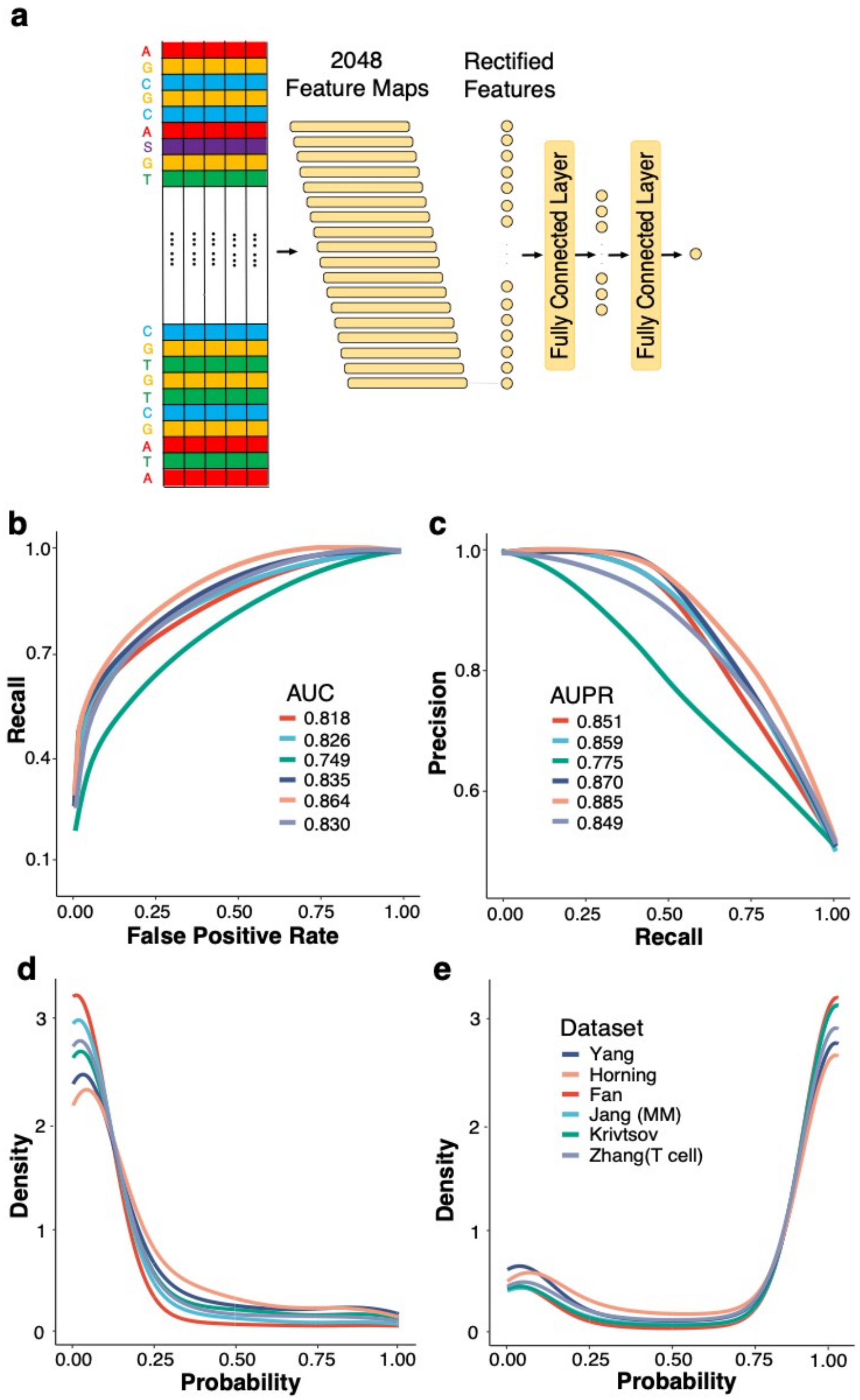


(**a**) The design of the CNN model (**b**) The ROCs of the CNN model for different single cell data sets. The AUCs are also shown. (**c**) The PR curves and their AUPRs of the CNN model. (**d**) The densities of the technical artefact score of gene fusions in the PCAWG study by the CNN models retrained using six different datasets. (**e**) The densities of the technical artefact score of chimeric reads. The models are retrained using different datasets.
